# Identification of epigenome-wide DNA methylation differences between carriers of *APOE* ε4 and *APOE* ε2

**DOI:** 10.1101/815035

**Authors:** Rosie M. Walker, Kadi Vaher, Mairead L. Bermingham, Stewart W. Morris, Andrew D. Bretherick, Yanni Zeng, Konrad Rawlik, Carmen Amador, Archie Campbell, Chris S. Haley, Caroline Hayward, David J. Porteous, Andrew M. McIntosh, Riccardo E. Marioni, Kathryn L. Evans

## Abstract

**BACKGROUND:** The *Apolipoprotein E* (*APOE*) ε4 allele is the strongest genetic risk factor for late onset Alzheimer’s disease, while the ε2 allele confers protection. Previous studies report differential DNA methylation of *APOE* between ε4 and ε2 carriers, but associations with epigenome-wide methylation have not previously been characterised.

**METHODS:** Using the EPIC array, we investigated epigenome-wide differences in whole blood DNA methylation patterns between Alzheimer’s disease-free *APOE* ε4 (n=2469) and ε2 (n=1118) carriers from the two largest single-cohort DNA methylation samples profiled to date. Using a discovery, replication and meta-analysis study design, methylation differences were identified using epigenome-wide association analysis and differentially methylated region (DMR) approaches. Results were explored using pathway and methylation quantitative trait loci (meQTL) analyses.

**RESULTS:** We obtained replicated evidence for DNA methylation differences in a ^~^169kb region, which encompasses part of *APOE* and several upstream genes. Meta-analytic approaches identified DNA methylation differences outside of *APOE:* differentially methylated positions were identified in *DHCR24, LDLR* and *ABCG1* (2.59 x 10^−100^≤*P*≤2.44 x 10^−8^) and DMRs were identified in *SREBF2* and *LDLR* (1.63 x 10^−4^≤*P*≤3.01 x 10^−2^). Pathway and meQTL analyses implicated lipid-related processes and high density lipoprotein cholesterol was identified as a partial mediator of the methylation differences in *ABCG1* and *DHCR24*.

**CONCLUSIONS:** *APOE* ε4 vs. ε2 carrier status is associated with epigenome-wide methylation differences in the blood. The loci identified are located in *trans* as well as *cis* to *APOE* and implicate genes involved in lipid homeostasis.

## 1. Background

The ε4 allele of the *apolipoprotein E* gene (*APOE*) is the strongest genetic risk factor for late-onset (>65 years) Alzheimer’s disease (AD) (1–3). Inheritance of one copy of this allele increases late-onset AD risk by two to four-fold, with two copies conferring an eight to twelve–fold increase in risk compared to the ε3/ε3 genotype (4, 5). The ε4 allele is also associated with a younger age-of-onset, with ε4 homozygotes having an average age-of-onset of 68 years compared to 84 years for ε3 homozygotes (4). In contrast, the ε2 allele has been associated with a ^~^50% reduction in AD risk compared to the ε3/ε3 genotype (5).

The three *APOE* alleles (ε2/ε3/ε4) are defined by two *APOE* exon 4 single nucleotide polymorphisms (SNPs) and encode functionally distinct ApoE isoforms. Isoform-dependent behaviours have been observed for many ApoE functions, including lipid metabolism, amyloid beta (Aβ) metabolism, tau phosphorylation, inflammation, and synaptic plasticity, with ApoE4 and ApoE2 conferring effects consistent with increased and reduced AD risk, respectively (6, 7).

Despite the wealth of evidence linking ApoE to processes implicated in AD pathogenesis, understanding of the specific mechanism(s) by which genetic variation at this locus alters risk remains incomplete. *APOE* genotype acts in conjunction with other genetic and/or environmental factors to confer AD risk: the lifetime risk of dementia or mild cognitive impairment is 31%-40% for ε4/ε4 homozygotes (8) but the effects of *APOE* ε4 have been shown to be modified by ethnic background and sex (5, 9). DNA methylation is associated with both genetic and environmental factors, and previous studies have identified associations with AD and neuropathological hallmarks of AD (10–12), AD risk factors (e.g. ageing (13), obesity (14) and lipid levels (15)), as well as modifiers of *APOE* genotype effects (e.g. sex (16) and ethnicity (17, 18)).

The two *APOE* haplotype-defining SNPs are located in a CpG island and have a direct effect on methylation by creating/destroying CpG sites (19). The *APOE* ε2/ε3/ε4 haplotype is associated with methylation at other CpG sites within *APOE* (20, 21) but, to date, associations with methylation across the epigenome have not been assessed. We hypothesised that characterising these associations would yield insights into the biological context in which *APOE* acts, thus facilitating the search for mechanisms conferring risk/resilience for AD. Importantly, by studying individuals who are free from AD, we have the potential to identify pathogenic processes that precede the onset of irreversible neurodegeneration.

## 2. Methods

### 2.1. Participants

The participants were selected from the Generation Scotland: Scottish Family Health Study (GS:SFHS) cohort (^~^24,000 participants aged ≥18 years at recruitment), which has been described previously (22, 23). The participants included in this study were of European (predominantly British) ancestry, following the exclusion of participants with likely recent Italian or African/Asian ancestry by principal components (PC) analysis (24). Participants attended a baseline clinical appointment at which they were phenotyped for social, demographic, health and lifestyle factors, completed cognitive assessments, and provided physical measurements and samples for DNA extraction. GS:SFHS obtained ethical approval from the NHS Tayside Committee on Medical Research Ethics, on behalf of the National Health Service (reference: 05/S1401/89) and has Research Tissue Bank Status (reference: 15/ES/0040).

### 2.2. Blood sample collection and DNA extraction

DNA was extracted from blood (9ml) collected in EDTA tubes using the Nucleon BACC3 Genomic DNA Extraction Kit (Fisher Scientific), following the manufacturer’s instructions (25).

### 2.3. Genotyping of *APOE*

The *APOE* ε2/ε3/ε4 haplotypes are defined by two SNPs, rs429358 and rs7412, which were genotyped using TaqMan probes at the Clinical Research Facility, Edinburgh.

### 2.4. Measurement of cholesterol levels

Total and high density lipoprotein (HDL) cholesterol were measured at the GS:SFHS baseline appointment and non-HDL cholesterol levels were calculated by subtracting HDL cholesterol from total cholesterol. The non-HDL cholesterol level reflects a combination of low density lipoprotein (LDL) cholesterol and very low-density lipoprotein.

### 2.5. Genome-wide DNA methylation profiling for EWAS analyses

DNA methylation was profiled using the Infinium MethylationEPIC BeadChip (Illumina Inc.) in a discovery (n=5190) and replication (n=4583) sample, as described previously (26–28) (Supplementary Methods). The discovery and replication samples were normalised separately and converted to M-values. The discovery data was corrected for relatedness (Supplementary Methods). Participants in the replication sample were unrelated (SNP-based relatedness<0.05) to each other and/or discovery sample participants.

Poor performing probes, X/Y chromosome probes and participants with unreliable self-report data or potential XXY genotype were excluded (Supplementary Methods). The final discovery dataset comprised M-values at 760,943 loci for 5087 participants; the replication dataset comprised M-values at 758,332 loci for 4450 participants. All subsequent analyses of the DNA methylation data were carried out using R versions 3.6.0., 3.6.1., or 3.6.2. (29, 30).

### 2.6. Statistical analyses

A flow chart indicating all analyses is presented in Figure 1.

**Figure 1.**
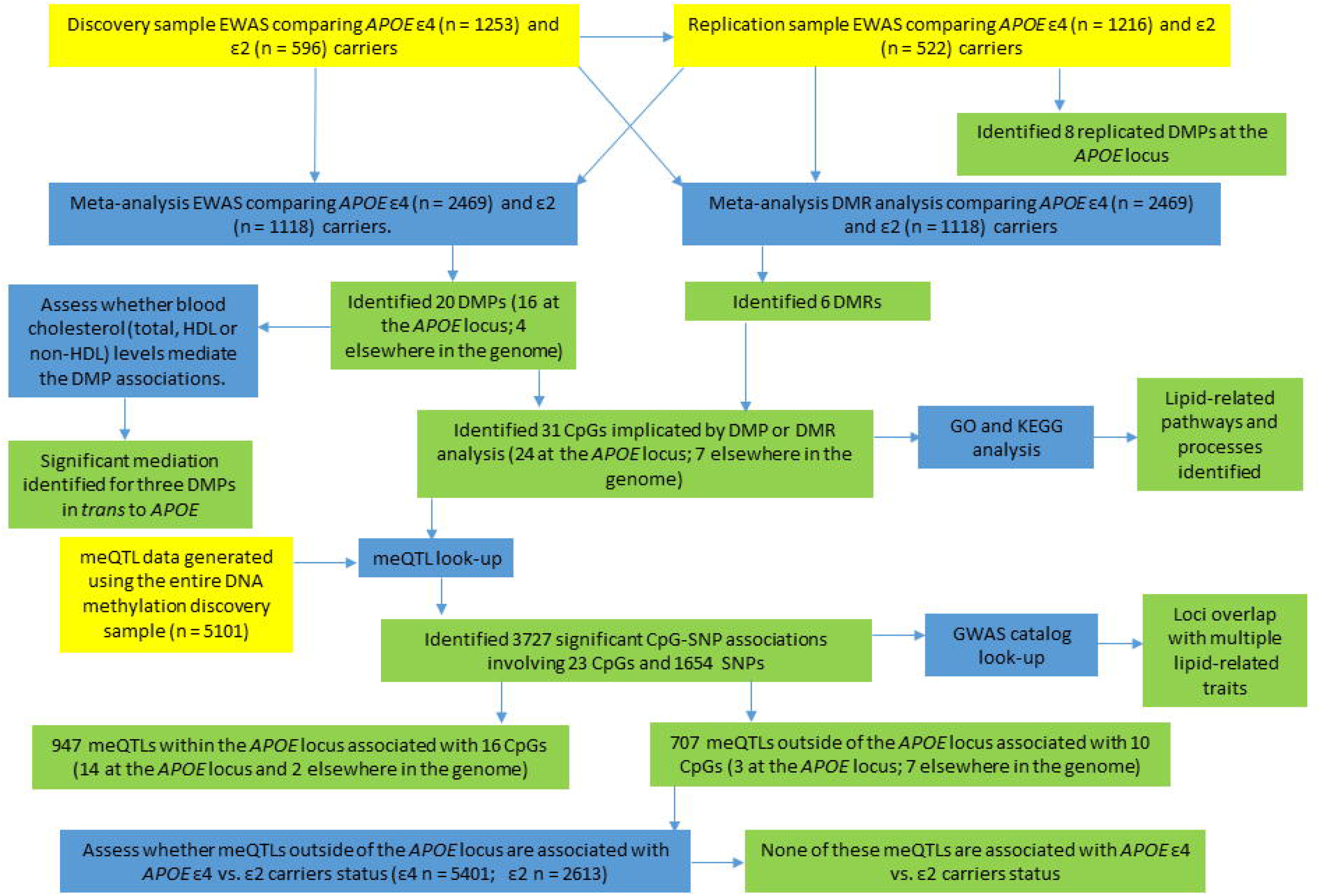
Flow chart indicating the analyses carried out in this study. Yellow boxes indicate datasets used for the analysis, blue boxes describe the analysis performed and green boxes contain the results of the analysis. Arrows indicate for which analyses the datasets were used, the order of the analyses and the results from each analysis.

### 2.7. Epigenome-wide association studies

EWASs were implemented using limma (31). CpG M-values were the dependent variable and *APOE* ε4 vs. ε2 carrier status (a binary variable indicating *APOE* ε4 carriers with a “1” and *APOE* ε2 with a “0”; ε4/ε2 and ε3/ε3 participants were excluded) was the predictor-of-interest. Participants self-reporting AD (n=five) were excluded. Additional covariates were included as below:

#### Discovery sample

CpG site (pre-corrected for relatedness, estimated cell counts and processing batch) ^~^ *APOE* ε4 vs. ε2 + age + sex + smoking status + pack years + 20 methylation PCs

#### Replication sample

CpG site (M-values) ^~^ *APOE* ε4 vs. ε2 + age + sex + smoking status + pack years + estimated cell counts (granulocytes, natural killer cells, B-lymphocytes, CD4+T-lymphocytes and CD8+T-lymphocytes) + processing batch + 20 methylation PCs

The variables “smoking status”, “pack years” and the methylation PCs are explained in the Supplementary Methods.

An additional sensitivity analysis of the replication sample was performed in which the first 10 genetic PCs, calculated using GCTA (32), were included. The decision to include 10 PCs was based on inspection of a scree plot (Supplementary Figure 1).

Limma was used to calculate empirical Bayes moderated t-statistics from which *P*-values were obtained. The significance threshold in the discovery sample was *P*≤3.6 x 10^−8^ (33). Sites attaining significance in the discovery sample were assessed in the replication sample using a Bonferroni-corrected threshold of 0.05/no. sites assessed.

### 2.8. EWAS meta-analysis

Inverse variance-weighted fixed effects meta-analyses of 756,971 sites common to the discovery and replication EWAS results were performed using METAL (34). Sites attaining a meta-analysis *P*≤3.6 x 10^−8^ were considered significant.

### 2.9. Comparison of DNA methylation levels between *APOE* haplotypes

For the differentially methylated positions (DMPs) identified through the EWAS meta-analysis, pairwise differences in methylation levels between carriers of the *APOE* ε2/ε2, ε2/ε3, ε3/ε3, ε3/ε4, and ε4/ε4 haplotypes in the discovery sample were investigated, using the R package lsmeans (35). *P*-values were adjusted using a Bonferroni correction to account for the 10 within-CpG comparisons performed for each of the 20 CpGs assessed (i.e. an adjustment was performed for 200 tests). Corrected *P*≤0.05 was considered statistically significant.

### 2.10. Identification of differentially methylated regions

DMRs associated with *APOE* ε4 vs. ε2 carrier status were identified using the dmrff.meta function from the dmrff R package (36). Putative DMRs were defined as regions containing two to thirty sites separated by ≤500 bp with EWAS meta-analysis *P*≤.05 and methylation changes in a consistent direction. Following dmrff’s subregion selection step, DMRs with Bonferroni-adjusted *P*≤.05 were declared significant.

### 2.11. Gene ontology/KEGG pathway analyses

Gene ontology (GO) and KEGG pathway analyses were implemented using missMethyl’s gometh function (37). The target list comprised probes that were suggestively associated with the phenotype-of-interest (*P*≤1 x 10^−5^) in the meta-EWAS or that contributed to a significant DMR (adjusted *P*≤0.05) and the gene universe included all analysed probes. Enrichment was assessed using a hypergeometric test, accounting for the bias arising from the variation in the number of probes-per-gene. Bonferroni-corrected significance thresholds of *P*≤2.21 x 10^−6^ and *P*≤1.48 x 10^−4^ were applied to account for the 22,578 GO terms and 337 KEGG pathways assessed.

### 2.12. Bootstrap mediation analysis

The roles of cholesterol levels (total cholesterol, HDL cholesterol and non-HDL cholesterol) in mediating any observed associations between *APOE* ε4 vs. ε2 carrier status and DNA methylation were assessed by bootstrap mediation analysis using the R package “mediation” (38). The analyses were performed using 10000 bootstrap samples in the discovery and replication samples separately and these results were then meta-analysed using inverse variance-weighted fixed effects meta-analyses to obtain meta-analyses *P*-values and effect estimates. Significant mediation was declared when the meta-analysis *P*-value met a Bonferroni-adjusted (to account for the assessment of 20 DMPs) significance threshold of *P*≤.05.

### 2.13. Genotyping and imputation

The genotyping and imputation of GS:SFHS to the Haplotype Reference Consortium reference panel release 1.1 (39) has been described previously (25, 40) (Supplementary Methods).

### 2.14. Identification of methylation quantitative trait loci

Methylation quantitative trait loci (meQTLs) were identified using the discovery sample. Following quality control, the data was normalised and corrected as described previously (41) (Supplementary Methods). Normalised and corrected data was available for 26 of the 31 CpGs-of-interest in this study. The resulting residuals were inverse rank normal transformed and entered as the dependent variable in simple linear model GWASs to identify meQTLs. GWASs were implemented using REGSCAN v0.5 (42). SNPs that were associated with a CpG with *P*≤1.92 x 10^−9^ (5 x 10^−8^/26) were declared to be meQTLs. SNPs located within one megabase up- or downstream of their associated CpG were defined as *cis* meQTLs; all other associated SNPs were defined as *trans* meQTLs. A look-up analysis of the GWAS catalog (43) (GWAS catalog v1.0.2., downloaded 07/09/20) was performed in which SNPs identified as meQTLs for the CpGs of interest were queried for their significant (*P*≤5 x 10^−8^) disease or trait associations in the GWAS catalog.

### 2.15. Association analyses of *APOE* ε4 vs. ε2 carrier status

Association analyses were performed to assess whether meQTLs for the meta-analysis DMPs are associated with *APOE* ε4 vs. ε2 carrier status and, therefore, might contribute to the differences in methylation observed between *APOE* ε4 and ε2 carriers. Association tests used BOLT-LMM (44) to perform linear mixed models in participants with available *APOE* genotypes (ε2 n=2613; ε4 n=5401). BOLT-LMM adjusts for population structure and relatedness between individuals whilst assessing association. Sex was included as a covariate. Associations were considered significant when *P*≤5 × 10^−8^.

## 3. Results

### 3.1. Sample demographics

The EWAS discovery sample comprised 1253 *APOE* ε4 and 596 *APOE* ε2 allele carriers and the replication sample comprised 1216 *APOE* ε4 and 522 *APOE* ε2 allele carriers. Twenty-seven ε2/ε2, 569 ε2/ε3, 2926 ε3/ε3, 1128 ε3/ε4 and 125 ε4/ε4 participants from the discovery sample were available for the pairwise analysis of genotypes. Key sample demographic information is presented in Supplementary Table 1.

### 3.2. Identification of differentially methylated positions and regions in *APOE* ε4 vs. ε2 carriers

An EWAS of *APOE* ε4 vs. ε2 carriers in the discovery sample identified eight significant DMPs, of which half were hypermethylated in *APOE* ε4 carriers. These DMPs had a mean absolute effect size of 0.070 (range: 0.033 – 0.103) and *P*-values ranging from 6.40 x 10^−56^ to 8.81 x 10^−9^. All eight sites were also significant (8.60 x 10^−49^≤*P*≤7.25 x 10^−6^) in the replication sample with a consistent direction of effect (mean absolute effect size = 0.102; range: 0.049 – 0.170; Supplementary Table 2). The eight sites are located in a ^~^169kb region on chromosome 19 (chr. 19: 45,242,346-45,411,802; GRCh37/hg19), which spans a region of the genome upstream of and including part of the *APOE* gene (chr19: 45,409,039-45,412,650; GRCh37/hg19). A sensitivity analysis of the discovery sample in which a methylation-based smoking score (45) was included as a covariate instead of the smoking covariates included in the original analysis (“smoking status” and “pack years”) produced highly similar results across all measured CpGs (correlation between effect sizes = 0.99, 95% confidence interval (CI): 0.99-0.99; *P*<2.2 x 10^−16^; Supplementary Table 2). An additional sensitivity analysis in which the first 10 genetic PCs were included as additional covariates in the analysis of the replication sample also produced results that were highly correlated with those from the original replication sample analysis (*r* = 1.00, 95% CI: 1.00-1.00, *P*<2.2 x 10^−16^; Supplementary Table 2).

Inverse variance-weighted fixed effects meta-analysis of the discovery and replication samples identified 20 DMPs, with *APOE* ε4 carrier status associated with hypomethylation at 13 (65%) of these sites. Across all 20 DMPs, the mean absolute effect size was 0.052 (range: 0.022 – 0.11) with *P*-values ranging from 2.80 x 10^−100^ to 2.4 x 10^−8^ (Table 1; Figure 2). Sixteen of these sites are located on chromosome 19q in a ^~^233kb region (chr19: 45,221,584 – 45,454,752; GRCh37/hg19) encompassing *APOE* and several surrounding genes (Supplementary Figure 2). Henceforth, the region containing *APOE* and neighbouring genes will be referred to as the “*APOE* locus”. The most significant DMP, cg13375295, is located ^~^4.5kb upstream of *Poliovirus Receptor-related 2 (PVRL2*), a gene situated ^~^16.5kb upstream of *APOE*. Four other DMPs (cg10762466, cg10178308, cg11643040 and cg06198803) are located either upstream or in the gene body of *PVRL2*. Two DMPs (cg06750524 and cg16471933) are located in *APOE:* cg06750524, the DMP with the largest effect size, in the intron between exons 2 and 3; and cg16471933 in exon 4, 139bp 5’ of rs429358, one of the *APOE* ε4/ε2-defining SNPs. Although both the *APOE* DMPs are more highly methylated in *APOE* ε4 carriers; the DMPs in the surrounding region do not show a consistent direction of effect.

**Table 1.** *APOE* ε4 vs. ε2-associated DMPs identified by meta-analysis of the discovery and replication EWASs

| Probe ID | Gene symbol | Gene<br>feature * | Chr. | BP<sup>†</sup> | Effect<sup>‡</sup> | SE | P value |
| --- | --- | --- | --- | --- | --- | --- | --- |
| cg13375295 |  |  | 19 | 45344725 | -0.1031 | 0.0049 | 2.80 x 10<sup>-100</sup> |
| cg06750524 | *APOE* | Body | 19 | 45409955 | 0.1122 | 0.008 | 1.05 x 10<sup>-44</sup> |
| cg16094954 | *BCL3* | TSS1500 | 19 | 45251180 | -0.0994 | 0.0081 | 8.18 x 10<sup>-35</sup> |
| cg10762466 |  |  | 19 | 45347693 | -0.0463 | 0.004 | 1.37 x 10<sup>-30</sup> |
| cg16471933 | *APOE* | Body | 19 | 45411802 | 0.0606 | 0.0055 | 7.17 x 10<sup>-28</sup> |
| cg10178308 | *PVRL2* | TSS200 | 19 | 45349383 | 0.1075 | 0.0103 | 2.04 x 10<sup>-25</sup> |
| cg27087650 | *BCL3* | Body | 19 | 45255796 | 0.0455 | 0.0044 | 3.77 x 10<sup>-25</sup> |
| cg04488858 |  |  | 19 | 45242346 | -0.0514 | 0.0065 | 2.25 x 10<sup>-15</sup> |
| cg11643040 | *PVRL2* | Body | 19 | 45361327 | -0.0278 | 0.0038 | 1.46 x 10<sup>-13</sup> |
| cg26631131 |  |  | 19 | 45240591 | 0.0298 | 0.0042 | 2.45 x 10<sup>-12</sup> |
| cg17901584 | *DHCR24;RP11-67L3.4* | TSS1500 | 1 | 55353706 | -0.0403 | 0.0058 | 3.58 x 10<sup>-12</sup> |
| cg06198803 | *PVRL2* | Body | 19 | 45371896 | -0.041 | 0.006 | 1.04 x 10<sup>-11</sup> |
| cg16740586 | *ABCG1* | Body | 21 | 43655919 | 0.0332 | 0.005 | 3.58 x 10<sup>-11</sup> |
| cg03793277 | *APOC1* | TSS1500 | 19 | 45416910 | -0.0304 | 0.0049 | 5.99 x 10<sup>-10</sup> |
| cg06500161 | *ABCG1* | Body | 21 | 43656587 | 0.0247 | 0.0042 | 2.67 x 10<sup>-9</sup> |
| cg09555818 | *APOC2;APOC4* | 5' UTR;<br>1<sup>st</sup> exon | 19 | 45449301 | -0.0531 | 0.0091 | 5.77 x 10<sup>-9</sup> |
| cg13119609 | *APOC2;APOC4* | 5' UTR;<br>1<sup>st</sup> exon | 19 | 45449297 | -0.0464 | 0.008 | 5.84 x 10<sup>-9</sup> |
| cg15233575 | | | 19 | 45221584 | -0.0223 | 0.0039 | $7.17 \times 10^{-9}$ |
| cg14645843 | | | 19 | 45454752 | -0.0346 | 0.0062 | $2.31 \times 10^{-8}$ |
| cg19751789 | *LDLR* | | 19 | 11199944 | -0.0338 | 0.0061 | $2.43 \times 10^{-8}$ |

Abbreviations: BP, base position; Chr., chromosome; SE, standard error; TSS, transcription start site;
UTR, untranslated region
\* Gene feature: 5' UTR: between the TSS and the ATG; Body: between the ATG and the stop codon;
TSS200: within 200 bases 5' of the TSS; TSS1500: within 1500 bases 5' of the TSS.
† Base position in genome assembly hg19/GRCh37
‡ Effect direction is relative to carriers of the $\epsilon 2$ allele

**Figure 2.**
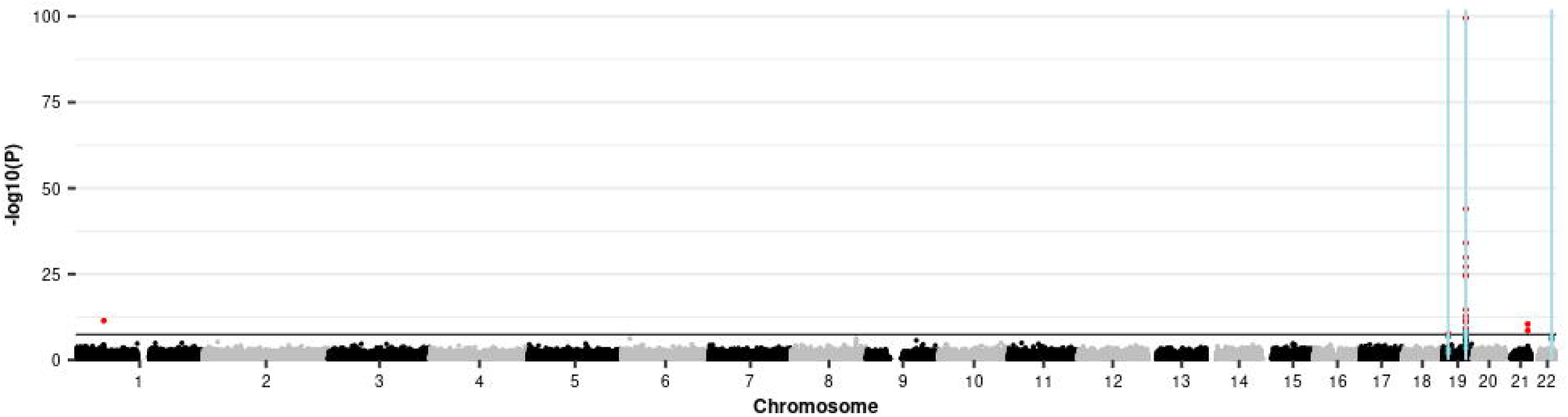
Manhattan plot showing the *APOE* ε4 vs. ε2 carrier EWAS and DMR meta-analyses results. Each point represents one of the 772,453 loci included in the EWAS meta-analysis, with the point’s position being determined by genomic position (x-axis) and significance in the EWAS metaanalysis (−log_10_ *P* value; y-axis). Sites attaining genome-wide significance (*P* ≤ 3.6 x 10^−8^) are indicated in red and those that are involved in a significant DMR (Bonferroni-correct *P* ≤ 0.05) are indicated in blue. The locations of DMRs are further indicated by vertical blue lines. The solid horizontal line is the threshold for genome-wide significance (*P* ≤ 3.6 x 10^−8^).

Four DMPs are located outside of chromosome 19q: cg17901584, 785bp upstream of the 24-*dehydrocholesterol reductase* (*DHCR24*) gene on chromosome 1; cg19751789, 94bp upstream of the *low density lipoprotein receptor* (*LDLR*) gene on chromosome 19p; and two, cg16740586 and cg06500161, are located 668bp apart in the same intron of multiple *ATP Binding Cassette Subfamily G Member 1* (*ABCG1*) isoforms.

To further investigate the pattern of methylation observed at these 20 DMPs, pairwise comparisons were performed between carriers of the following *APOE* haplotypes: ε2/ε2, ε2/ε3, ε3/ε3, ε3/ε4, and ε4/ε4. These analyses revealed a range of allele-associated methylation patterns, which are depicted in Supplementary Figure 3 and described in Supplementary Table 3. Carriers of the *APOE* ε2 allele (ε2/ε2 or ε2/ε3) differed from ε3/ε3 homozygotes at 14 of the DMPs, whilst carriers of the *APOE* ε4 allele (ε4/ε4 or ε3/ε4) differed from ε3/ε3 homozygotes at four DMPs. Dosage effects were observed at two DMPs for ε2 carriers (Supplementary Figure 3 A and S) and one DMP for ε4 carriers (Supplementary Figure 3B), although the small numbers of participants who are homozygous for *APOE* ε2 (n = 27) and ε4 (n = 128) likely rendered our study underpowered to detect all dosage effects. For the two DMPs located within the *APOE* gene (cg06750524 and cg16471933), an increase in mean methylation levels was observed from ε2/ε2 homozygotes to ε3/ε3 homozygotes, with a further increase to the ε4/ε4 group (Supplementary Figure 3B and E). At the four DMPs outside of the *APOE* locus, the methylation differences appear to be predominantly driven by the ε2 allele (Supplementary Figure 3K, M, O and S).

Differentially methylated regions (DMRs) were identified using a meta-analysis approach, which identified six significant regions (Supplementary Figure 4). Across all the DMRs, the mean absolute effect size was 0.182 (range: 0.135 – 0.231) and Bonferroni-adjusted *P*-values ranged from 1.63 x 10^−4^ to 3.01 x 10^−2^ (Table 2).Three of the DMRs are located at the *APOE* locus, two are in the first intron of *Sterol Regulatory Element Binding Transcription Factor 2* (*SREBF2*) on chromosome 22, and the other is in the putative promoter of *LDLR* on chromosome 19p. All but one of the DMRs, which is located 190 bp upstream of the *apolipoprotein C1 pseudogene 1* (*APOC1P1*) at the *APOE* locus, are hypomethylated in *APOE* ε4 carriers. Only one of the DMRs, located in an exon of a read-through transcript involving *apolipoprotein C2* (*APOC2*) and *apolipoprotein C4* (*APOC4*), contains CpGs that were identified as DMPs (cg13119609 and cg09555818).

**Table 2.** Significant DMRs identified through DMR meta-analysis of the discovery and replication sample EWAS results

| Chr. | Coordinates<sup>*</sup> | Gene symbol | Effect<sup>†</sup> | SE | Adj. *P* value<sup>‡</sup> | CpGs |
| --- | --- | --- | --- | --- | --- | --- |
| 19 | 45449297-45449301 | *APOC2;APOC4* | -0.231 | 0.0364 | 1.63 x 10<sup>-4</sup> | cg13119609;<br>cg09555818 |
| 19 | 45449099-45449150 | *APOC4-*<br>*APOC2;APOC2;APOC4* | -0.212 | 0.0356 | 0.00203 | cg01958934;<br>cg10872931 |
| 19 | 11199851-11199903 | *LDLR* | -0.135 | 0.0245 | 0.0290 | cg07960944;<br>cg05249393;<br>cg22381454; |
| 22 | 42230879-42230899 | *SREBF2* | -0.189 | 0.0329 | 0.00755 | cg15128785;<br>cg12403973 |
| 22 | 42229983-42230138 | *SREBF2* | -0.176 | 0.0312 | 0.0118 | cg09978077;<br>cg16000331 |
| 19 | 45429771-45429870 | *APOC1P1* | 0.148 | 0.0269 | 0.0301 | cg23184690;<br>cg08121984 |

Abbreviations: Chr., chromosome; SE, standard error; Adj., adjusted; CpGs, cytosine and guanine nucleotides linked by a phosphate bond
<sup>\*</sup> DMR start and end coordinates in genome assembly hg19/GRCh37
<sup>†</sup> Effect direction is relative to carriers of the ε2 allele
<sup>‡</sup> Bonferroni-adjusted *P* value

GO analysis was carried out using the 19 Entrez IDs mapping to the 46 CpG sites with a meta-EWAS **P*≤1* x 10^−5^ or that contributed to a significant DMR. This identified 78 significant GO terms (Table 3; Supplementary Table 4), the most significant of which was “cholesterol metabolic process” (*P*=2.00 x 10^−11^). Significant enrichment for the KEGG pathways “cholesterol metabolism” (*P*=5.93 x 10^−10^) and “steroid biosynthesis” (*P*=1.22 x 10^−4^) was also observed.

**Table 3.**
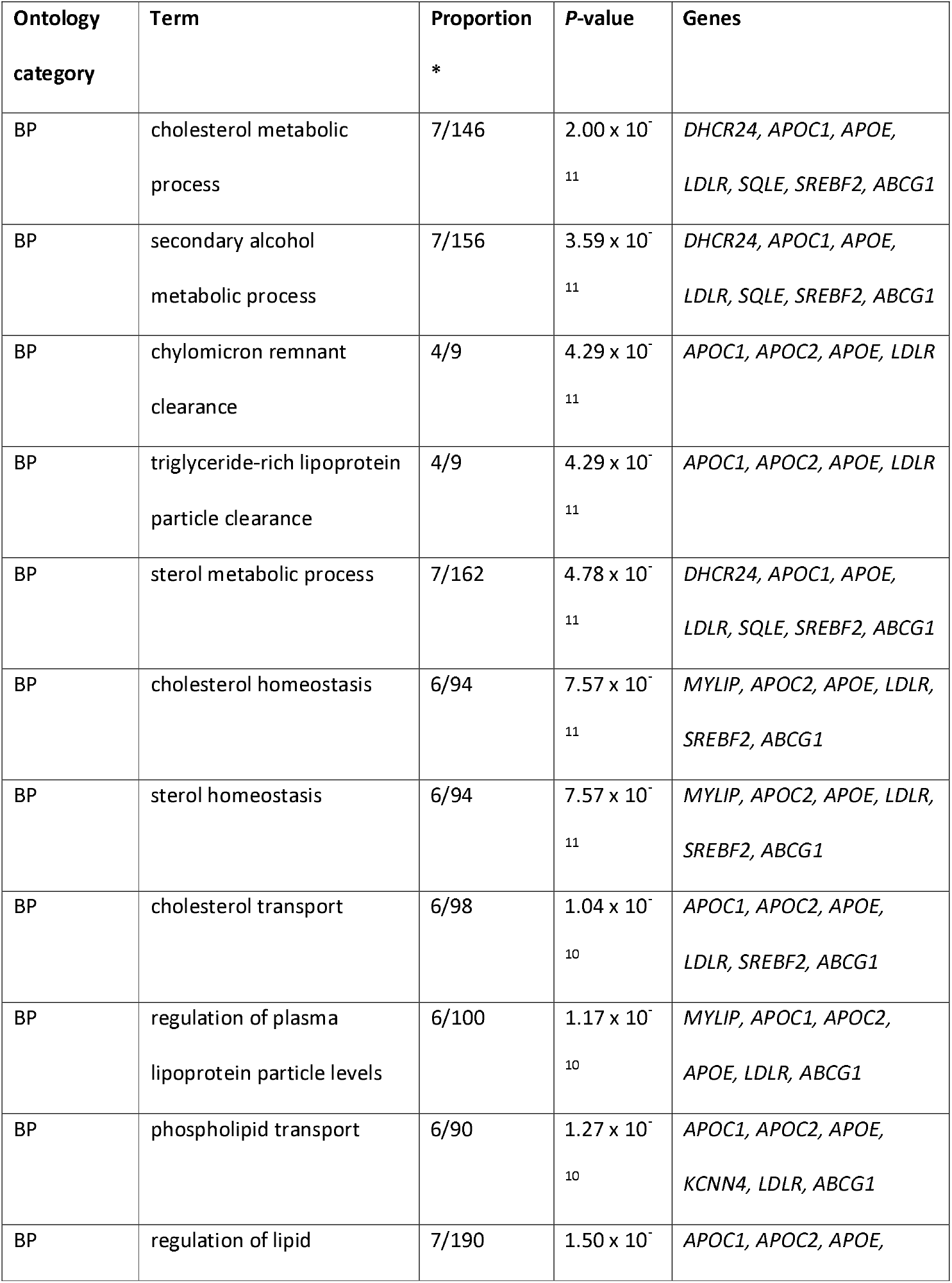
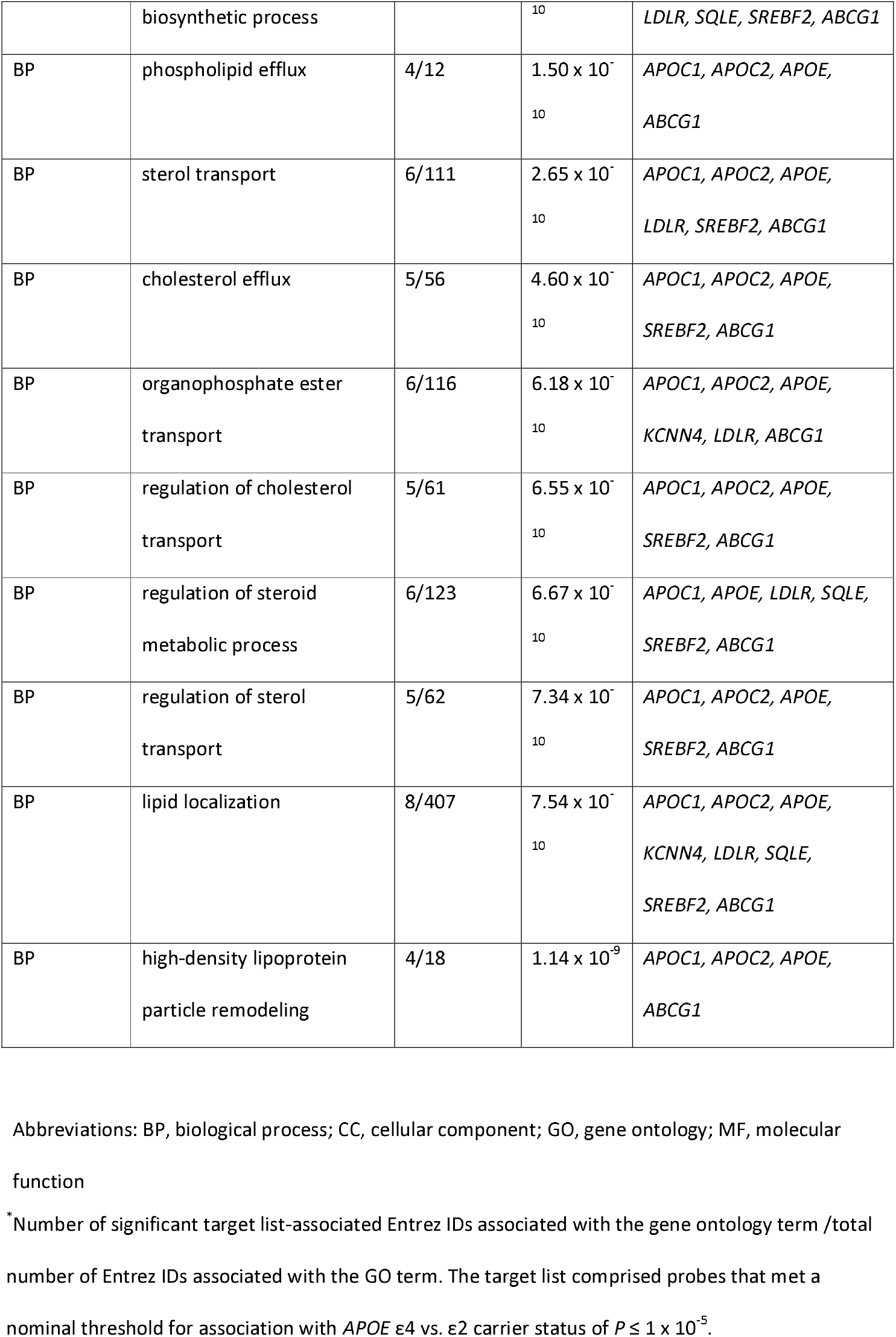
Top 20 GO terms showing significant enrichment for *APOE* ε4 vs. ε2-associated differentially methylated loci

### 3.3. Assessment of the role of cholesterol in mediating methylation differences between *APOE* ε4 and ε2 carriers

Given the well-established role of ApoE in cholesterol metabolism (6), bootstrap mediation analyses were performed to assess the role of cholesterol levels (total, HDL or non-HDL cholesterol) in mediating the association between *APOE* ε4 vs. ε2 carrier status and methylation at the 20 meta-analysis DMPs. Inverse variance-weighted fixed effects meta-analysis of the bootstrap mediation analyses in the discovery and replication samples identified HDL cholesterol as a significant mediator of the associations with the two *ABCG1* DMPs cg06500161 (effect size = 0.006; effect size standard error = 0.001; *P*=1.18 x 10^−6^) and cg16740586 (effect size = 0.004; effect size standard error = 0.001; *P*=4.93 x 10^−5^), and the *DHCR24* promoter DMP, cg17901584 (effect size = −0.007; effect size standard error = 0.001; *P*=6.04 x 10^−6^), for which it mediated 25.2%, 11.5%, and 18.2% of the relationship, respectively (Supplementary Table 5). For some sites, inspection of the *P*-values indicated total and non-HDL cholesterol to be significant mediators but the proportion of the relationship between *APOE* ε4 vs. ε2 carrier status and methylation attributable to the mediator was negative (Supplementary Table 5). This indicates that, at these sites, the direction of the association between the cholesterol phenotypes and methylation is the opposite to the direction of the association between *APOE* ε4 vs. ε2 carrier status and methylation.

### 3.4. Assessment of meQTLs associated with loci that are differentially methylated between *APOE* ε4 and ε2 carriers

To explore the DMP and DMR CpGs further, meQTL analyses were performed. Whilst it was expected that meQTLs for the DMP and DMR CpGs would be identified at the *APOE* locus, the identification of meQTLs outside of this locus would be of particular interest. Should meQTLs outside of the *APOE* locus be found to be show non-random segregation with *APOE* ε4 vs. ε2 carrier status, these meQTL SNPs might contribute to the methylation differences observed in this study and *APOE* genotype effects more generally.

It was possible to assess meQTLs for 26 of the 31 CpGs of interest (from the DMP and DMR analyses); amongst these CpGs, 23 were associated with a meQTL. In total, 3727 significant CpG-SNP associations were identified for the 23 CpGs, involving 1654 unique SNPs (Figure 3; Supplementary Table 6). Unsurprisingly, more than half of the meQTLs (n=947) were located in a ^~^719kb region (chr19: 45,004,645− 45,723,446; GRCh37/hg19) spanning *APOE*. The *APOE* region meQTLs are associated with 16 CpGs, of which 14 are located at the *APOE* locus. None of these meQTLs is associated with all 16 CpGs: two are each associated with nine CpGs: rs7412, one of the *APOE* ε2/ε3/ε4-defining SNPs; and rs41290120, an intronic *PVRL2* SNP that is in high linkage disequilibrium with rs7412 with D’ = 0.85 in the British population (46). The two CpGs associated in *trans* with SNPs in the *APOE* region are cg16000331 in *SREBF2* and cg19751789 in *LDLR*.

**Figure 3.**
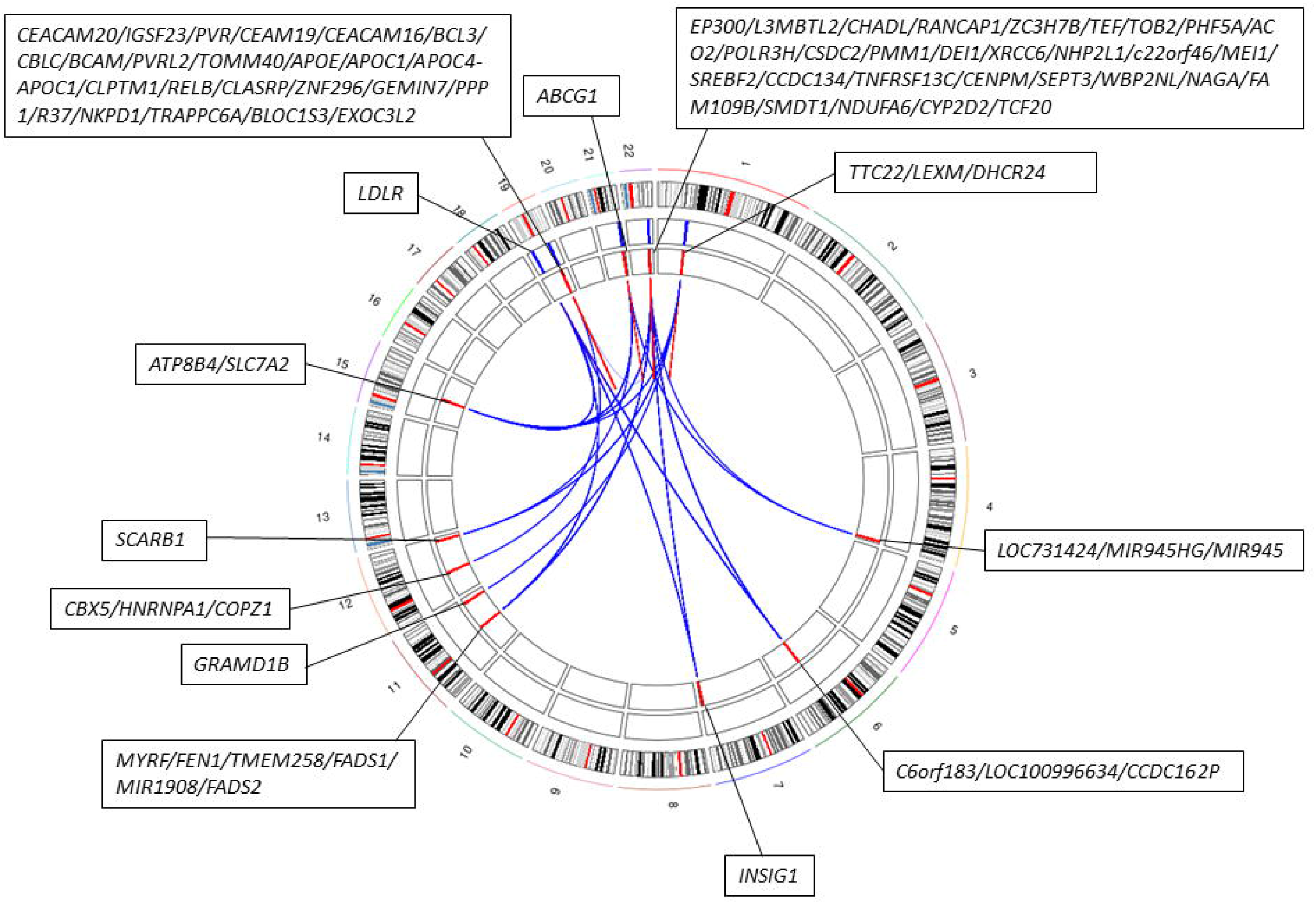
Circular plot indicating the locations of *APOE* ε4 vs. ε2 carrier-associated DMP and DMR CpGs. The first track shows a chromosome ideogram (hg19/GRCh37). The genomic locations of CpGs identified as being DMPs or in DMRs identified in *APOE* ε4 vs. ε2 carriers are indicated by blue lines on the second track and the meQTLs associated with these CpGs are indicated by the red lines on the third track. The connections between CpGs and meQTLs indicate regulatory relationships (*cis* interactions in red; *trans* interactions in blue). Gene symbols for genes located in each CpG/meQTL-harbouring region are indicated.

Outside of the *APOE* locus, the remaining 707 meQTLs, which are associated with 10 CpGs, are located in 11 genomic regions (Figure 3; Supplementary Table 7), with each region containing meQTLs associated with between one and eight CpGs-of-interest. To assess whether these meQTLs might contribute to *APOE* ε4 vs. ε2-associated methylation differences, their association with *APOE* ε4 vs. ε2 carrier status was assessed. No significant associations were observed, suggesting that the *APOE* ε4 vs. ε2-associated methylation differences are predominantly driven by genotype at the *APOE* locus.

To investigate potential trait/disease associations with variation in methylation levels at the CpGs-of-interest, the GWAS catalog was queried(43). This identified 234/1654 meQTLs as having genome-wide significant associations with 316 traits (Supplementary Table 8). More than one third of the associations are with a lipid-related traits, including LDL, HDL and total cholesterol levels. As expected, many of the meQTL SNPs within the *APOE* locus have previously been associated with AD and related traits, such as “Cerebrospinal fluid p-tau levels”, “Cerebral amyloid deposition (PET imaging)” and “Cognitive decline”. Interestingly, five SNPs located outside of the *APOE* locus have also been associated with traits related cognitive ability (“Cognitive ability, years of educational attainment or schizophrenia (pleiotropy) “, “General cognitive ability”, “Intelligence” and “Self-reported math ability”). Four of these SNPs encompass the 3’ end of *CCDC134* and most of the neighbouring *SREBF2*. Between them, these four SNPs are associated in *cis* with methylation at the four CpGs forming the two *SREBF2* DMRs. The fifth SNP, which is located on chromosome 6 in the pseudogene *CCDC162P*, is associated with methylation at CpGs in *SREBF2* and *LDLR*. Three meQTL SNPs have been associated with several age-related disorders (e.g. heart failure, stroke, and cancer) and endophenotypes of these disorders (including cholesterol levels, blood pressure and blood glucose) in a pleiotropic GWAS meta-analysis (47).

## 4. Discussion

We performed the first epigenome-wide comparison of DNA methylation between carriers of the *APOE* ε4 and ε2 haplotypes, which confer risk for and protection from AD, respectively. In large discovery and replication samples, we confirm the presence of *APOE* haplotype-associated methylation differences in *APOE*, demonstrate that differences in methylation at the *APOE* locus span a broad genomic locus encompassing several genes, and find evidence for altered methylation at sites unlinked to the *APOE* locus. The observed methylation differences are located in a network of genes involved in lipid metabolism and homeostasis.

Methylation differences were identified using discovery, replication and meta-analysis EWASs and DMR analysis. Eight DMPs located on chromosome 19 in a ^~^169kb region spanning from upstream of *BCL3* to the *APOE’s* fourth exon showed replicated association. An additional twelve DMPs, eight of which are located in a ^~^233kb region at the *APOE* locus, were identified by meta-analysing the discovery and replication samples. DMR analysis identified six regions of differential methylation, both within and outside of the *APOE* locus.

Within the *APOE* gene, two DMPs, cg06750524, in the second intron, and cg16471933, in the fourth exon, were identified. *APOE* ε4 carriers showed higher methylation levels at both. This observation directly replicates a previous study (21) and is in line with Foraker et al.’s observation of increased methylation of the *APOE* exon 4 CpG island in ε4 carriers (20). Moreover, we have previously demonstrated (48) that the pattern of methylation in *APOE* in our sample is consistent with that described by Ma et al. (2015) (21). Pairwise comparisons revealed differences in *APOE* methylation to be driven both by differences between ε2 carriers and ε3/ε3 homozygotes and ε4 carriers and ε3/ε3 homozygotes. One interpretation of this observation is that the spectrum of methylation at the *APOE* DMPs reflects the spectrum of AD risk conferred by different AD genotypes. It is clear, however, that additional, likely experimental, studies are required to assess the implications of the observed methylation pattern.

The differentially methylated CpGs at the *APOE* locus span a broad region that encompasses several genes containing AD-associated variants (49). Long-ranging linkage disequilibrium in the region complicates the interpretation of association signals; however, conditional analysis and fine-mapping studies suggest the presence of multiple independent AD risk loci across the region (3, 49). As such, the methylation differences observed in this study may be associated with variants that, whilst being in LD with the *APOE* ε2/ε4-defining SNPs, confer risk via different pathways to these SNPs. This notion is supported by the observation that SNPs that define an *APOE* ε4-independent AD-risk haplotype in *PVRL2* (49) are highly significant meQTLs for the most significant DMP identified in this study.

Beyond the *APOE* locus, DMPs were identified in an *ABCG1* intron, and upstream of *DHCR24* and *LDLR*. Comparisons with ε3/ε3 homozygotes suggested the ε2 allele to be the primary driver of these differences, suggesting the possibility that altered methylation of genes involved in lipid metabolism might contribute to this allele’s protective effects. DMRs were identified in the gene body of *SREBF2* and in the putative promoter region of *LDLR*. The CpGs involved in the DMPs and DMRs located outside of the *APOE* locus are associated with several meQTLs, with all of the CpGs except those involved in the *LDLR* DMR being associated with meQTLs in *cis* as well as in *trans*. Our findings did not, however, support a role for *cis* meQTLs for these CpGs driving associations with *APOE* ε4 vs. ε2 carrier status.

The genes outside of the *APOE* locus that harbour differentially methylated CpGs are implicated in lipid metabolism or homeostasis. *ABCG1*, which is highly expressed in the brain, encodes a cholesterol and phospholipid transporter and is involved in regulating the sterol biosynthetic pathway (50). *DHCR24*, which encodes the cholesterol biosynthesis enzyme 3ß-hydroxysterol-Δ24 reductase, also known as seladin-1, plays a neuroprotective role in AD-related stress conditions, including Aβ toxicity, oxidative stress and inflammation (51, 52). The alteration of seladin-1 expression in mouse brain and human neuroblastoma cell cultures has been shown to affect β-secretase processing of amyloid precursor protein, with reduced seladin-1 being associated with an increased rate of Aβ production (53). Future studies should assess whether methylation-associated differences in the brain expression of seladin-1 (6) might mediate the established associations between *APOE* ε4 vs. ε2 haplotype and Aβ production. The *LDLR* gene encodes the LDL receptor, one of the neuronal receptors capable of mediating the endocytosis of ApoE, thus, maintaining brain cholesterol homeostasis. *LDLR* expression is regulated, in part, by *SREBF2*, a transcriptional regulator of sterol-regulated genes, which contains a SNP that is associated both with *SREBF2* expression and CSF levels of the AD biomarkers Aβ and tau (54).

The link between *APOE* ε4 vs. ε2-associated methylation differences and lipid-related processes and pathways was further supported by GO and KEGG analyses, the identification of meQTLs for the differentially methylated CpGs, which were clustered in genomic regions that contain several lipid-related genes, and their GWAS-associated phenotypes. It would be of interest to investigate the mechanisms underlying the clustering of meQTLs in these genomic regions. Future studies might assess, for example, the extent to which meQTLs associated with the differentially methylated CpGs are enriched in these regions and whether they disproportionately affect certain sequence motifs. Previous EWASs have also identified associations between some of the *APOE* ε4 vs. ε2-associated CpGs and cholesterol levels: the *DHCR24* (cg17901584), *ABCG1* (cg06500161) and *SREBF2* (cg16000331) DMPs have been associated with HDL cholesterol, total cholesterol and triglyceride levels (15, 55–57). Comparisons with previous EWASs are, however, limited by the fact that the majority of previous EWASs used the 450K array, which, does not contain 10 of the *APOE* ε4 vs. ε2-associated CpGs.

As differences in lipid metabolism between carriers of the *APOE* ε4 and ε2 haplotypes are well-documented (6), we assessed whether variation in blood cholesterol levels might mediate the observed *APOE* ε4 vs. ε2-associated methylation differences. HDL cholesterol was found to be a partial mediator of the relationship between *APOE* ε4 vs. ε2 carrier status and methylation at three loci located outside of the *APOE* locus (two within *ABCG1* and one in the promoter of *DHCR24*), thus suggesting one mechanism that might underlie these *trans* effects. Consistent with our observation that methylation differences at these loci appear to be predominantly driven by *APOE* ε2 carriers (when compared to *APOE* ε3/ε3 homozygotes), higher HDL cholesterol levels have been reported in carriers *of APOE* ε2 (58). The effect of HDL cholesterol on methylation varied between the three loci, with *APOE* ε2 carriers showing increased methylation at the site located in the *DHCR24* promoter and decreased methylation at the two *ABCG1* sites. This suggests that increased HDL cholesterol levels do not exert a general effect on methylation but rather that methylation varies in a locus-specific manner in response to variation in HDL levels. It should be noted that an assumption of this analysis is that reverse causation does not exist between the outcome, methylation, and the mediator, cholesterol. Previous Mendelian Randomisation studies have predominantly supported this premise (59, 60); however, the ability to identify robust genetic instruments has limited both the number of methylation sites assessed and the ability to assess reverse causation. Limitations to the GS:SFHS cholesterol data should also be noted when interpreting these findings: triglyceride levels were not measured, preventing LDL cholesterol assessment; and blood samples were not taken at a consistent time of day or after fasting.

The cross-sectional nature of this study precludes the observed methylation differences being interpreted as conferring risk, protection or compensation. Comparison of methylation at these loci in *APOE* ε4 and ε2 carriers with AD would be useful in addressing this question; however, the optimum study design would involve the longitudinal assessment of the trajectory of ε4 vs. ε2-associated methylation differences in AD-free individuals in midlife who either do or do not later develop AD. These analyses are currently not feasible due to the small sizes of existing AD patient blood-based DNA methylation samples and insufficient follow-up time of large population-based samples.

Studies assessing the association of neuropathological hallmarks (neuritic plaque burden and/or neurofibrillary tangles) of AD with DNA methylation in the brain have not identified the loci identified in the present study (10, 12, 61). Although the phenotypes assessed differ, the existence of *APOE* haplotype-associated differences in Aβ metabolism and tau phosphorylation (6) suggest that some degree of overlap might be expected. The neuropathological hallmarks of AD are, however, complex phenotypes and *APOE* haplotype will be one of many contributing factors (De Jager et al. (10) reported that *APOE* ε4 could account for 13.9% of the variance in NP burden observed in their participants). In addition, the smaller samples assessed by De Jager et al. (10), Lunnon et al. (12) and Smith et al. (61) may have been inadequately powered to detect any methylation differences driven by *APOE* haplotype. Differences in age and methylation profiling platform are also likely to limit comparability: the participants assessed in these studies were much older (mean age >75 years) than those assessed in our study (mean age ^~^ 50 years) and array differences mean that only two thirds of our DMP/DMR probes were assessed. Two important corollaries of the age difference are that brain-based studies are more likely to (i) suffer from survivor bias and (ii) be better suited to investigating end-of-disease processes. It is also important to note that *APOE* is involved in multiple processes, with *APOE* ε4 conferring risk for AD, at least in part, via mechanisms that are not related to Aβ or tau pathology. A recent study has indicated that *APOE* ε4-associated breakdown of the blood-brain barrier in the hippocampus and medial temporal lobe contributes to *APOE* ε4-associated cognitive decline independently of Aβ or tau (62).

The blood provides an easily accessible tissue that can be repeatedly sampled to characterise pre-morbid markers of risk. The extent to which it can provide mechanistic insights into diseases that are considered predominantly brain-based, however, is a perennial subject of debate. *Cis* meQTL effects tend to be highly correlated (*r* = 0.78) between the blood and the brain (63), supporting the use of the blood to study the effects of genetic risk factors for brain-based diseases. It is also important to note the increasing recognition of the role of peripheral processes in conferring risk for AD (64). As the blood provides a conduit by which many circulating factors (e.g. plasma proteins and microbial metabolites) reach the brain and affect brain ageing (65), assessing DNA methylation in the blood is likely to be informative regarding systemic factors contributing to AD pathogenesis. Although *APOE* is synthesised separately in the blood and the brain and neither *APOE* nor cholesterol can cross the blood-brain barrier (66, 67), there is cross-talk between brain and blood cholesterol via oxysterols (67), levels of which vary by *APOE* ε2/ε3/ε4 haplotype (68). Peripheral hypercholesterolemia has been associated with increased oxysterol levels in the brain, which have been implicated in with production and accumulation of Aβ, increased neuroinflammation and neuronal death (67).

The association between *APOE* genotype and AD varies between populations (5), with studies in populations of Hispanic and African ancestry often reporting attenuated effect sizes for the ε4 allele compared to studies involving European and Asian participants (69, 70). Moreover, Rajabli et al. (70) have shown that genetic variation local to *APOE* is likely to confer protection from the effects of the ε4 allele in individuals of African ancestry. As the participants in the present study were of European ancestry, it should be noted that these findings are likely to be European-specific and future studies should assess their generalisability and relevance to AD pathogenesis in other populations.

## 5. Conclusions

This is the first study to characterise epigenome-wide DNA methylation differences between carriers *of APOE* ε4 and ε2. In AD-free individuals, we identified several methylation differences both at the *APOE* locus and in the rest of the genome, which converge on lipid-related pathways. Strengths of the study include the large samples available for EWAS analysis, the epigenome-wide approach, the use of a well-phenotyped cohort with genotype data, and the avoidance of reverse causation by studying AD-free participants. Future studies should investigate the causal relationship between *APOE* genotype, DNA methylation and lipid-related processes and their role in AD pathogenesis.

## Abbreviations

Aβ: amyloid beta
AD: Alzheimer’s disease
CI: confidence interval
DMP: differentially methylated position
DMR: differentially methylated region
EWAS: epigenome-wide association study
GS:SFHS: Generation Scotland: Scottish Family Health Study
HDL: high density lipoprotein
LDL: low density lipoprotein
PC: principal component
SNP: single nucleotide polymorphism

## Declarations

### Ethics approval and consent to participate

GS:SFHS obtained ethical approval from the NHS Tayside Committee on Medical Research Ethics, on behalf of the National Health Service (reference: 05/S1401/89) and has Research Tissue Bank Status (reference: 15/ES/0040), providing generic ethical approval for a wide range of uses within medical research. All experimental methods were in accordance with the Helsinki declaration.

### Consent for publication

Not applicable

### Availability of data and materials

According to the terms of consent for GS:SFHS, access to data must be reviewed by the GS Access Committee.

### Authors’ contributions

Conception and design: RMW, KLE, REM; data analysis: RMW, KV, MLB, ADB, CH. Drafting the article: RMW and KLE; data preparation: RMW, MLB, SWM, KR, AC, ADB, YZ, CA; data collection: AMM, KLE, CSH, DJP. Revision of the article: RMW, KV, MLB, ADB, YZ, CA, AC, CSH, DJP, REM, KLE; all authors read and approved the final manuscript.

## Acknowledgements

We are grateful to all the families who took part, the general practitioners and the Scottish School of Primary Care for their help in recruiting them, and the whole Generation Scotland team, which includes interviewers, computer and laboratory technicians, clerical workers, research scientists, volunteers, managers, receptionists, healthcare assistants and nurses.

