## Supplementary Methods for "Identification of epigenome-wide DNA methylation differences between carriers of *APOE* ε4 and *APOE* ε2"

1. **Genome-wide DNA methylation profiling**

Whole blood genomic DNA (500ng) from 9785 participants was treated with sodium bisulphite using the EZ-96 DNA Methylation Kit (Zymo Research, Irvine, California), following the manufacturer’s instructions. DNA methylation was profiled using the Infinium MethylationEPIC BeadChip (Illumina Inc.), according to the manufacturer’s protocol. DNA methylation was profiled in the discovery (n = 5200) and replication (n = 4585) samples at separate time points and quality control and normalisation of the samples was carried out separately. R version 3.3.2 [1] was used for the discovery sample and 3.4.3 [2] for the replication sample. These steps have been described in detail previously [3-5]; however, briefly, poor-performing probes (discovery sample N = 5910; replication sample N = 8878) and participants (discovery sample N = 80; replication sample N = 123), together with participants for whom there was a mismatch between their predicted sex (based on DNA methylation data) and their recorded sex (discovery sample N = 12; replication sample N = 12), were excluded from both samples. A further seven participants were excluded from the discovery sample as they were identified as being genetic outliers based on principal components analysis of genetic data [6]. Prior to use in all downstream analyses except the meQTL analyses, additional participants were excluded from the discovery sample: 10 participants whose DNA samples were subsequently discovered to have come from saliva; three participants who had answered “yes” for all self-reported conditions and one whose methylation data indicated likely XXY genotype.

Probes that had been predicted to have off-target effects or contain a SNP in the final five 3’ bases or at the site of single base extension for type I probes [7, 8] were excluded (N = 84,352). Probes on the X (N = 19,090) or Y (N = 537) chromosomes were also excluded.

The final discovery sample dataset comprised corrected M-values at 760,943 loci measured in 5087 participants (5101 for meQTL analyses), while the replication sample dataset comprised M-values at 758,332 loci measured in 4450 participants. Subsets of the discovery (n = 1839) and replication (n = 1738) samples, selected by *APOE* genotype, were assessed in the epigenome-wide association studies (EWASs). The pairwise comparison of APOE genotypes was carried out using the discovery sample and involved 27 ε2/ε2, 569 ε2/ε3, 2926 ε3/ε3, 1128 ε3/ε4 and 125 ε4/ε4 participants. All subsequent analyses of the DNA methylation data were carried out using R versions 3.6.0., 3.6.1. or 3.6.2. [9].

2. **Additional pre-processing steps prior to EWAS analyses**

The discovery and replication samples were normalised (separately) using the dasen method from the wateRmelon R package [10] and converted to M-values using the beta2m function in lumi [11].

As the discovery sample included related participants, the M-values for CpGs on autosomal chromosomes in this sample were pre-corrected for relatedness, estimated white blood cell proportions and processing batch using DISSECT [12]. This was achieved by saving the residuals from a mixed linear model that included methylation as the dependent variable and the following predictor variables: a genetic relatedness matrix fitted in a leave-one-chromosome-out fashion (i.e. SNPs on the same chromosome as the CpG were excluded); proportions of granulocytes, natural killer cells, B-lymphocytes, CD4+ T-lymphocytes and CD8+ T-lymphocytes estimated using an implementation of Houseman et al.’s [13] cell type prediction algorithm in minfi [14]; and a variable that indicated the batch in which array hybridisation, staining and scanning took place. Participants in the replication sample were selected to be unrelated (SNP-based genetic relatedness < 0.05) to each other and/or participants in the discovery sample.

3. **Smoking-related variables**

“Smoking status” is a categorical variable with five levels (current smoker, former smoker who gave up less than 12 months ago, former smoker who gave up 12 months or more ago, never smoked, undeclared) and “pack years” is a measure that indicates an individual’s lifetime exposure to tobacco. Pack years were calculated by multiplying the years an individual had smoked for by the maximum number of packs of cigarettes they ever smoked per day (a pack = 20 cigarettes). A conversion was used for cigars (a cigar = four cigarettes) and rolling tobacco (a 25g pack = 50 cigarettes).

4. **Methylation principal components**

Methylation principal components (PCs) were calculated from M-values that had been pre-corrected for age, sex, estimated cell counts, processing batch and relatedness (discovery sample only) using the R package FactoMineR [15]. Prior to inclusion as covariates, associations between the PCs and *APOE* ε4 vs. ε2 status in the discovery and replication samples were assessed by two-tailed independent samples t-tests. After applying a Bonferroni correction to account for the 20 tests performed, no PC was associated with *APOE* ε4 vs. ε2 status (min. *P* = 0.74).

5. **Genotyping and imputation**

The genotyping of GS:SFHS has been described in detail previously [16, 17]. Briefly, genome-wide genotype data was generated using either the Illumina Human OmniExpressExome-8-v1.0 BeadChip or the Illumina HumanOmniExpressExome-8 v1.2 BeadChip by the Genetics Core Laboratory at the Clinical Research Facility, Edinburgh, Scotland (www.wtcrf.ed.ac.uk). Genotype data was processed using the IlluminaGenomeStudio Analysis software v.2011.1 (Illumina, San Diego, CA). Quality control was carried out to remove SNPs with: <98% call rate; a Hardy-Weinberg equilibrium *P* ≤ 1 x 10^-6^; or a MAF < 1%. Imputation was then performed on 602,450 (561,125 following the MAF filtering step used for GWAS) autosomal SNPs following the Sanger Imputation Service pipeline (<https://imputation.sanger.ac.uk/>), which uses the Haplotype Reference Consortium reference panel release 1.1 [18]. Imputed SNPs with an info score ≥ 0.6 and MAF ≥ 0.01 were used for meQTL analyses, while imputed SNPs with an info score ≥ 0.4 and MAF ≥ 0.05 were used for association analysis.

6. **Identification of methylation quantitative trait loci**

MeQTLs were identified using the entire discovery sample. Following quality control, the data was normalised and corrected as described previously [19]. Briefly, normalisation was carried out using preprocessNoob in the R package minfi [14] and linear mixed modelling was used in two stages to produce corrected data, as follows:

Step 1: M-values ~ random(A + P) + fix(centre + batch + year + Sentrix position + top 50 control probe PCs)

The random effect “A” is a matrix representing appointment date and location in which elements representing individuals visiting the same clinic at the same date were set to one, diagonal elements were set to one, and other off-diagonals were set to zero. The random effect “P” is a matrix representing Sentrix slide in which elements representing individuals with the same Sentrix slide ID were set to1, diagonals set to one, and other off-diagonals were set to zero. The fixed effects are as follows: “centre” indicates the clinic in which the blood sample was taken; “batch” indicates the methylation profiling batch; “year” indicates the year in which the participant visited the clinic; “Sentrix position” indicates the position of the array on the Sentrix slide; and “top 50 control probe PCs” are the first 50 PCs calculated using the control probes included on the EPIC array.

Step 2: residual M-values from step 1 ~ random(G + K + F + S + C) + fix(sex + season + appointment day of week + age + age^2^ + estimated cell proportions (granulocytes, B-lymphocytes, natural killer cells, CD4+ T-lymphocytes, and CD8+ T-lymphocytes) + appointment time)

The random effects, “G”, “K”, “F”, “S”, and “C”, are a genomic relationship matrix, a kinship relationship matrix, and three environmental relationship matrices (full sibling relationships, couple relationships and nuclear family relationships). The fixed effect “season” indicates the season of the participants’ appointment, and estimated cell proportions for granulocytes, B-lymphocytes, natural killer cells, CD4+ T-lymphocytes, and CD8+ T-lymphocytes were generated using an implementation of Houseman et al.’s [13] cell type prediction algorithm in minfi [14]. Following the removal of probes that have been predicted to bind sub-optimally [7, 8], corrected data was available for 573,027 sites, including 26 of the 31 sites-of-interest in this study. The resulting residuals were inverse rank normal transformed before being entered as the dependent variable in simple linear model GWASs (implemented using REGSCAN v0.5 [20]) to identify meQTLs. SNPs that were associated with a CpG with *P* ≤ 1.85 x 10^-9^ (5 x 10^-8^/27) were declared to be meQTLs.
