## Supplementary material legends for "Identification of epigenome-wide DNA methylation differences between carriers of *APOE* ε4 and *APOE* ε2"

**Supplementary Table 1.** Sample demographic information for the discovery and replication samples and the individual *APOE* genotype groups within the discovery sample. The following information is shown: mean age; percentage of female participants; percentages of current smokers, former smokers who quit less than a year ago, former smokers who quit a year or more ago, never smokers and those that didn’t declare their smoking status; mean pack years (amongst current and former smokers).

**Supplementary Table 2.** Differentially methylated positions (DMPs) showing replicated association with *APOE* ε4 vs. ε4 carrier status and sensitivity analyses results. DMPs identified in the discovery sample (*P* ≤ 3.6 x 10^-8^) that showed association (*P* ≤ .00625 (0.05/8)) with a consistent direction of effect in the replication sample were deemed to have replicated. The probe ID, symbol of the gene in which the probe maps (if any), probe location (CHR and MAPINFO; hg19/GRCh37), genomic features associated with the probe (Body: within the gene body; TSS200: within 200 bp of the transcription start site; TSS1500: within 1500 bp of the transcription start site), beta coefficient, standard error of the beta, and *P*-value in the discovery (dis) and replication (rep) samples are shown for each replicated DMP. A sensitivity analysis of the discovery sample was performed in which a methylation based smoking-score (Maas et al., 2019) was included as a covariate in place of the “smoking status” and “pack years” covariates used in the original analysis (dis_Maas). A sensitivity analysis of the replication sample was performed in which ten genetic principal components were included as additional covariates (rep_10_genetic_PCs).

**Supplementary Table 3.** Pairwise comparison of DNA methylation levels between *APOE* genotypes for each of the 20 CpGs identified as being a DMP in the meta-analysis of the discovery and replication samples. The probe ID, contrast, effect size (estimate), standard error of the effect size (SE), t-statistic, and Bonferroni-adjusted *P*-value (adj. *p*-value) are shown for each CpG.

**Supplementary Table 4.** Gene ontology (GO) terms showing significant enrichment (*P*≤2.21 x 10^-6^) for the CpG sites identified as showing suggestive (*P*≤1 x 10^-5^) association with *APOE* ε4 vs. ε2 carrier status through DMP analysis or which contributed to a significant DMR. GO terms are categorised as referring to biological processes (BP), cellular compartments (CC) or molecular functions (MF). The “N” column indicates the total number of Entrez IDs associated with the GO term, the “DE” column indicates the number of significant target list-associated Entrez IDs associated with the gene ontology term, and the genes columns indicate the genes containing suggestively significant CpG sites that map to each term.

**Supplementary Table 5.** Assessment of total, HDL or non-HDL cholesterol as potential mediators of *APOE* ε4 vs. ε2 carrier status on methylation. For each DMP identified in the meta-analysis of the discovery and replication samples, the potential roles of total, HDL or non-HDL cholesterol levels as mediators of the observed associations with methylation were assessed by bootstrap mediation analysis (n = 10000 bootstraps) in the discovery and replication samples and the results meta-analysed. For each DMP, the probe ID, symbol of the gene in which the probe maps (if any), probe location (CHR and MAPINFO; hg19/GRCh37), direction of the effect (+: more methylation in *APOE* ε4 carriers; -: less methylation in *APOE* ε4 carriers) and *P-*value for the original analysis (without co-varying for cholesterol) are shown together with bootstrap effect estimates (Mediator effect), standard errors (Mediator SE), *P-*values (Mediator *P*), Bonferroni-adjusted *p*-values (Mediator *P.adj*) and the proportion of the total effect accounted for by the mediator (Mediator prop) for each of total, HDL and non-HDL cholesterol. Bold indicates significant mediation.

**Supplementary Table 6.** meQTLs for the *APOE* ε4 vs. ε2-associated DMPs and DMR sites. For each meQTL, the RS ID (rsid), SNP ID (chr_BP), SNP location (SNP_chr and SNP_pos; hg19/GRCh37), the assessed allele (a1), other allele (a0), the frequency of a1, the beta coefficient, standard error of the beta, *P-*value, variance explained in the residualised phenotype (r2), the associated CpG’s probe ID, the CpG location (CpG_chr and CpG_pos; hg19/GRCh37), and the type of meQTL (cis or trans) are shown

**Supplementary Table 7.** meQTLs grouped by genomic region. The meQTLs associated with the *APOE* ε4 vs. ε2-associated DMPs and DMRs map to 12 genomic regions. For each region, the location of the region (Chr. and start and end coordinates; hg19/GRCh37), its size, the total number of meQTLs , the number of *cis* and *trans* meQTLs, and the number of *APOE* ε4 vs. ε4-associated CpGs are shown.

**Supplementary Table 8.** GWAS catalog entries for the meQTLs associated with the *APOE* ε4 vs. ε2-associated DMPs and DMR sites. The GWAS catalog (<https://www.ebi.ac.uk/gwas/>) was queried for each of the SNPs that is an meQTL for the loci showing differential methylation between *APOE* ε4 and ε2 carriers and associations meeting a threshold of *P* ≤ 5 x 10^-8^ are shown. For each association, the SNP, location of the SNP (Chr and Pos; hg19/GRCh37), the symbol of the gene in which the SNP maps (if any; NR = not recorded), the associated trait, *P-*value, and a link to the study in which association was detected are shown.

**Supplementary Figure 1**. Scree plot showing the eigenvalues of the first 50 genetic principal components for the Generation Scotland: Scottish Family Health Study.

**Supplementary Figure 2.** The genomic region encompassing *APOE*, which contains 16 of the *APOE* ε4 vs. ε2-associated DMPs identified by meta-analysis (chr19: 45,221,584 – 45,454,752; GRCh37/hg19). The upper panel, downloaded from the UCSC Genome Browser (GRCh37/hg19) indicates the location of genes (NCBI RefSeq gene track). The lower panel indicates CpG sites in the region. The position of the CpG on the y-axis is determined by the –log_10_ of its meta-analysis *P*-value and its position on the x-axis is determined by its genomic location. The dashed red line indicates the threshold for significance (*p* ≤3.6 x 10^-8^) and CpGs attaining significance are labelled with their identifier. The two *APOE* SNPs, rs429358 and rs7412, which define the *APOE* ε2/ε 3/ε4 haplotype are also indicated.

**Supplementary Figure 3.** Bar charts showing the mean methylation levels (M-values) in the discovery sample for the 20 meta-DMPs split by *APOE* haplotype (ε2/ε2, ε2/ε3, ε3/ε3, ε3/ε4, and ε4/ε4. Error bars indicated standard errors of the mean. Pairwise differences were assessed using Tukey’s test of Honestly Significant Differences and significant differences are indicates by asterisks (**P* ≤ .05, ***P* ≤ .005, ****P* ≤ .0001).

**Supplementary Figure 4.** The genomic regions containing the six identified DMRs. **A**. indicates the three DMRs within the *APOE* locus (i. cg01958934 and cg10872931; ii. cg01958934 and cg10872931; iii. cg23184690 and cg08121984); **B**. shows the *LDLR* DMR; and **C**. shows the two *SREBF2* DMRs (i. cg15128785 and cg1240397; ii. cg09978077 and cg16000331). Each plot shows an extended region spanning from 10 kb downstream of the most 5’ DMR CpG to 10 kb upstream of the most 3’ DMR CpG. In each plot, the upper panel, downloaded from the UCSC Genome Browser (GRCh37/hg19) indicates the location of genes (NCBI RefSeq gene track). The lower panel indicates CpG sites in the region. The position of the CpG on the y-axis is determined by the –log_10_ of its meta-analysis *P*-value and its position on the x-axis is determined by its genomic location. The dashed red line indicates the threshold for significance (*p* ≤3.6 x 10^-8^) in the EWAS. The vertical blue lines indicate the locations of CpGs that are involved in a DMR and these CpGs are labelled with their probe ID.
