## Supplementary figures and images for "Identification of epigenome-wide DNA methylation differences between carriers of *APOE* ε4 and *APOE* ε2"

### Supplementary Figure 1

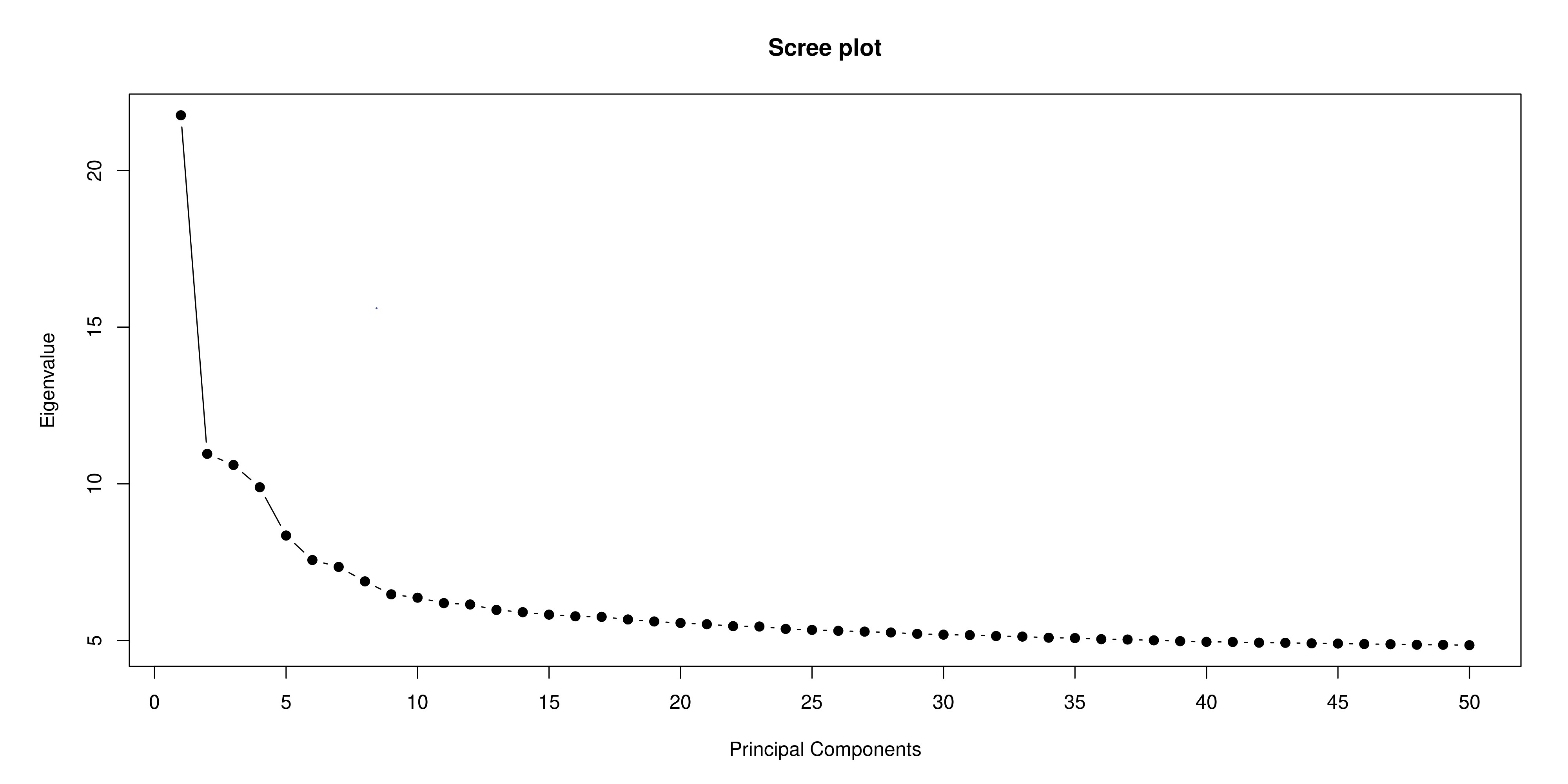

### Supplementary Figure 2

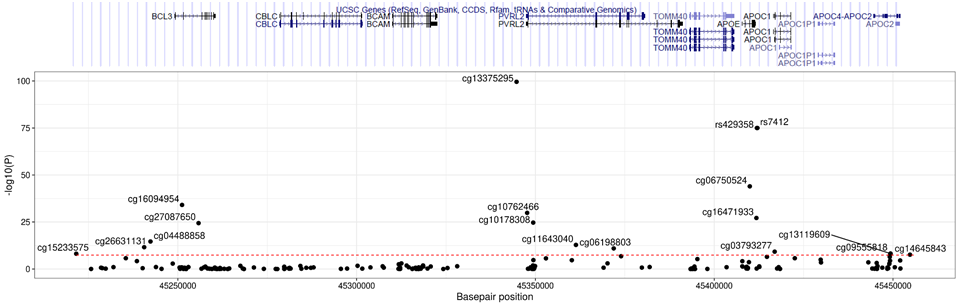

### Supplementary Figure 3

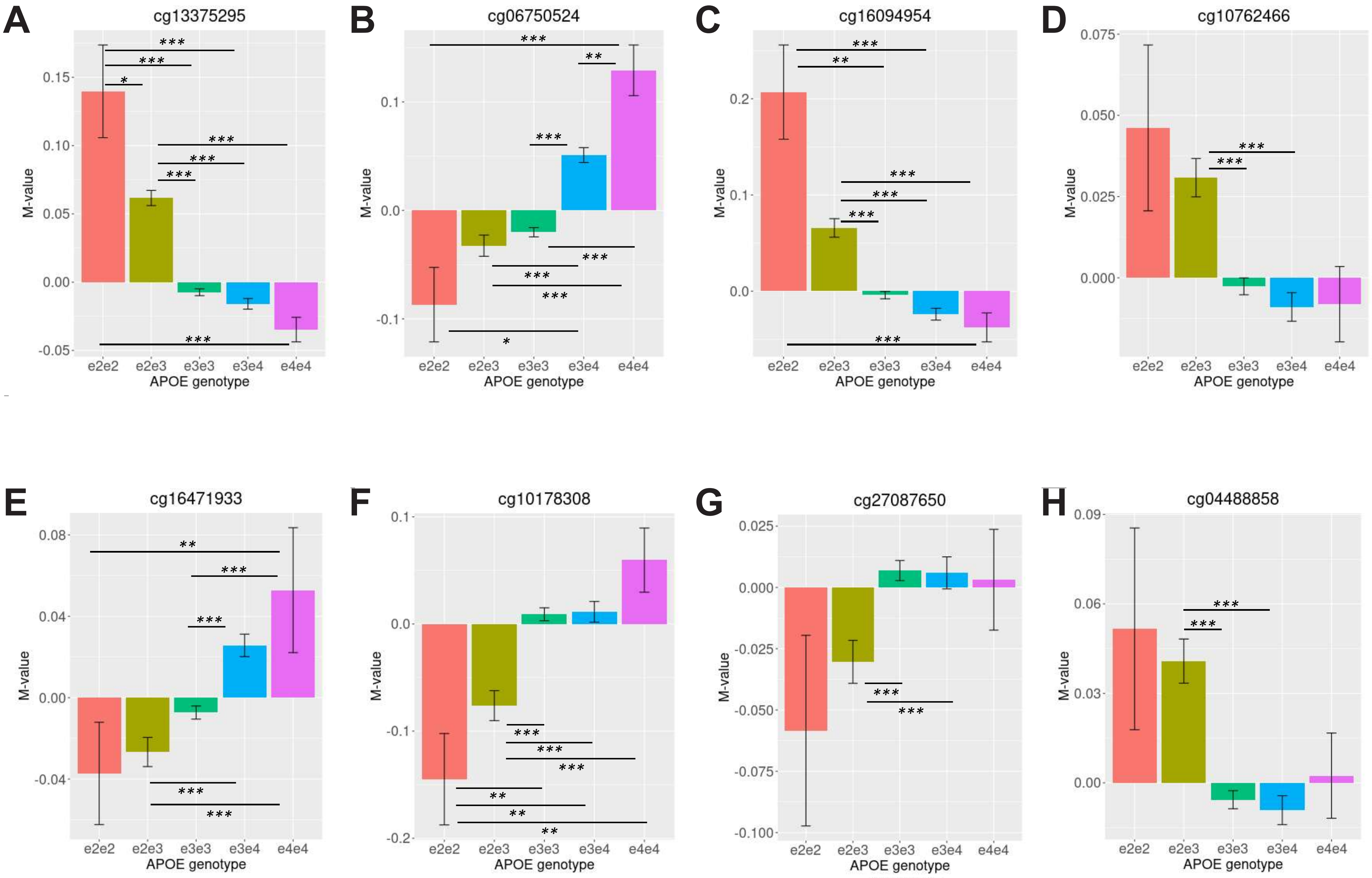

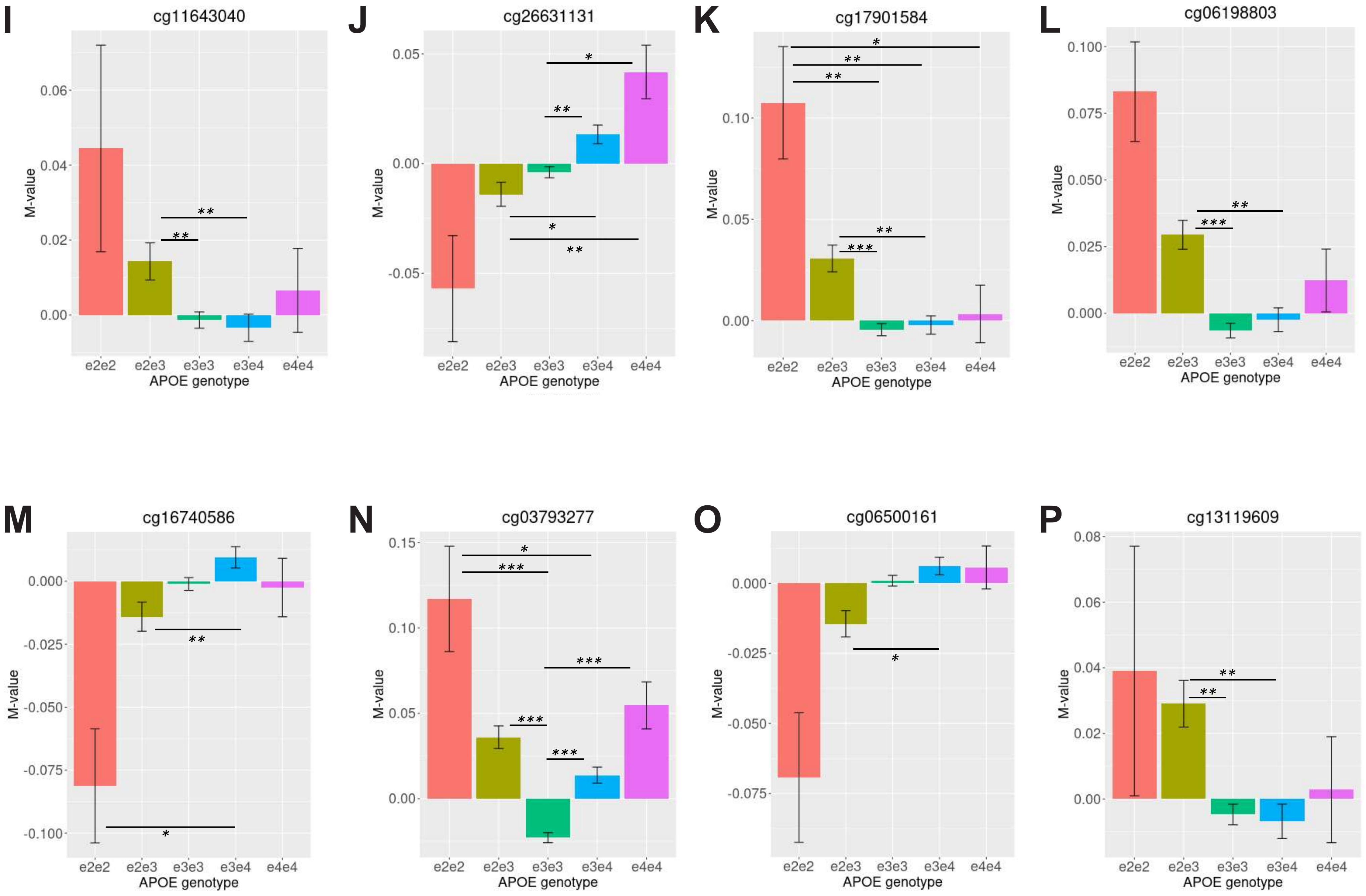

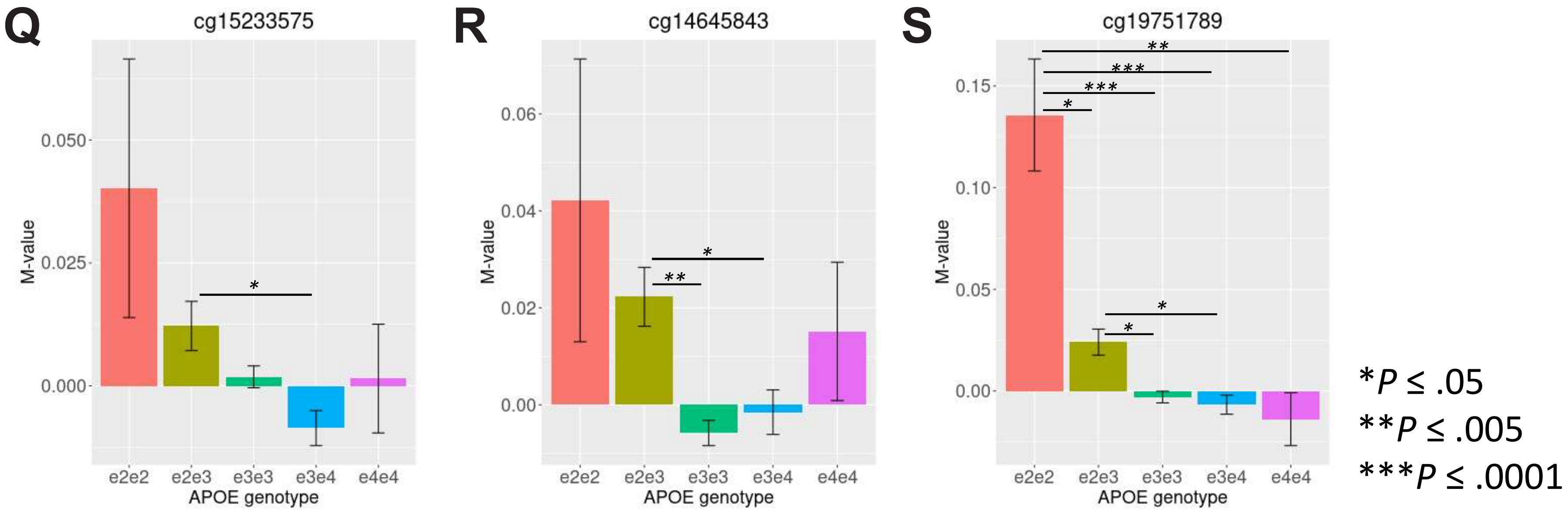

### Supplementary Figure 4

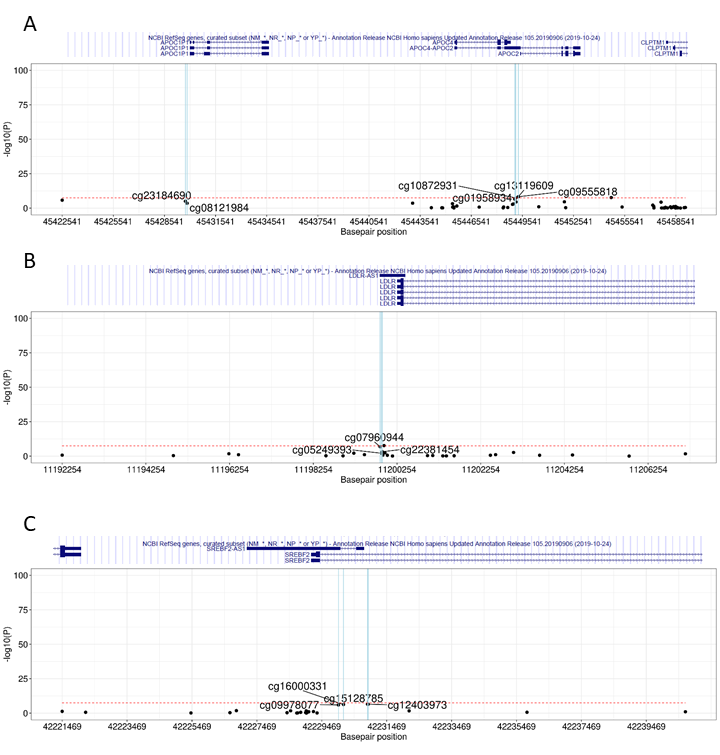
